# Characterising the spatio-temporal threats, conservation hotspots, and conservation gaps for the most extinction-prone bird family (Aves: Rallidae)

**DOI:** 10.1101/2021.01.20.427508

**Authors:** Lucile Lévêque, Jessie C. Buettel, Scott Carver, Barry W. Brook

## Abstract

With thousands of vertebrate species now threatened with extinction, there is an urgent need to understand and mitigate the causes of wildlife collapse. As distinct evolutionary clades can follow different routes to endangerment, there is value in taxon-specific analyses when assessing species’ vulnerability to threats and identifying gaps in conservation actions. Rails (Aves: Rallidae), being the most extinction-prone bird Family globally, and with one third of extant rail species now threatened or near-threatened, are an emphatic case in point. Yet even for this well-studied group, there is uncertainty in our understanding of what factors might be causing this vulnerability, whether the current threats are consistent with those that led to recent extinctions, and ultimately, what conservation actions might be necessary to mitigate further losses. Here, we undertook a global synthesis of the temporal and spatial threat patterns for Rallidae and determined conservation priorities and gaps. We found two key pathways in the threat pattern for rails. One follows the same trajectory as extinct rails, where island endemic and flightless rails are most threatened, mainly due to invasive predators. The second, created by the recent diversification of anthropogenic activities, involves continental rails (generally in the Neotropics), threatened most commonly by agriculture, natural-system modifications and residential and commercial development. Conservation efforts around most-at-risk species should be adapted according to the most relevant geographic scale (bioregions or countries), and principal locality type of the population (continental or island endemic). Indonesia, the U.S.A., the United Kingdom, New Zealand, and Cuba were the priority countries identified by our classification system incorporating species’ unique evolutionary features and level of endangerment, but also among the countries that lack conservation actions the most. Future efforts should predominantly target improvements in ecosystem protection and management, as well as ongoing research and monitoring. Forecasting the impacts of climate change on island endemic rails and disentangling the specific roles of extrinsic and intrinsic traits (like flightlessness), will be particularly valuable avenues of research for improving our forecasts of rail vulnerability.

## INTRODUCTION

Up to a million species are now threatened with extinction, with many more predicted within the next decade (IPBES, 2019). Along with the unprecedent decline in avifauna abundance, including in once common species (e.g. in North America, Rosenberg *et al*., 2019) the need to understand and mitigate the causes of these biotic contractions is urgent if we wish to avert a major loss of species and ecosystem integrity this century. As the vulnerability to different threats varies across taxa (e.g., between different bird families; Owens & Bennett, 2000), there is value in taxon-specific analyses to inform conservation policy and decision making. This has been done for several bird groups including ducks, geese and swans (Green, 1996), sea birds (Croxall *et al*., 2012; Spatz *et al*., 2014; Dias *et al*., 2019) and parrots (Olah *et al*., 2016), but gaps exist for many bird taxa, including the rail family.

Rails (Aves: Rallidae) are a widespread bird family, ecologically diverse and inhabiting most habitats worldwide. They are also the most extinction-prone bird family, with 54 to 92% of all species going extinct after their first contact with humans during the mid-Holocene (representing between 200 and 2,000 estimated extinct species; Steadman, 1995; Curnutt & Pimm, 2001) They went through a second wave of extinction starting in the 16^th^ century when European settlers spread worldwide. In this recent extinction wave, they were once again the most represented bird family in the extinct species assemblage (15%, 24 species; BirdLife International, 2018). Five rails have gone extinct in the last 100 years (*Hypotaenidia pacifica, Porphyrio paepae, Zapornia palmeri, Hypotaenidia wakensis*, and *Hypotaenidia poeciloptera*) and the most recent extinction event was recognised in 1994 (IUCN, 2019).

As the magnitude, diversity and global reach of anthropogenic activities increase, species can be impacted by an amplified range of pressures and stressors – the IUCN Red List now references 38 different types of threats (IUCN, 2019). As an already extinction-prone bird family, Rallidae could be particularly vulnerable to contemporary threats and continue on their trajectory to extinction. Of the 144 recognised extant species of Rallidae, 13 are near-threatened and 35 are threatened (Vulnerable, Endangered or Critically endangered; IUCN, 2019). To date, our understanding of historical threats and causes of extinctions in rails has been limited and their pattern of endangerment has not been clearly characterised either spatially (where extinct/threatened species are distributed) or temporally (with respect to changes in threat patterns between extinct and contemporary species). Here we provide the first such descriptive assessment for Rallidae. We have three main aims: i) characterise the pattern of threats (historical and modern), focusing on four different spatial scales (globally, on continents, for island endemics and between bioregions); ii) track the time course of species’ conservation status (IUCN status) and gaps in conservation programs, and iii) identify ‘conservation hotspots’ for rails by creating a ranking system using rails’ conservation values and level of endangerment, so as to identify priority countries and bioregions.

## METHODS

### Database compilation

We compiled a database for all 144 species of extant rails (including 42 island endemic species) using the 2019 version of the International Union for Conservation of Nature (IUCN) Red List (IUCN, 2019) and the ‘Guide to the Rails, Crakes, Gallinules and Coots of the world’ (Taylor & van Perlo, 1998). We used the taxonomic classification followed by the IUCN which included the rallid family of Sarothruridae (some authors consider it separate from the Rallidae family, see Garcia-R. *et al*., 2014; Garcia-R *et al*., 2020). The Sarothruridae family contains 15 species, including two threatened species, and is mostly present in the Afrotropics. In preliminary analyses we evaluated the effect of excluding this group, finding no major effect on results or their interpretations (except a minor effect of Threat rank of countries, see Table S1) and, thus, elected to retain the Sarothruridae in the Rallidae family. Species considered ‘Data deficient’ were excluded from the analysis (Table S2). Similarly, the New Caledonian rail (*Gallirallus lafresnayanus*) and the Samoan moorhen (*Pareudiastes pacificus*) are two ‘Critically endangered’ rail species that have not been seen with certainty since the 19^th^ century and are suspected to be extinct (IUCN, 2019); they were also excluded from our analysis. We considered island endemic species as those restricted to one or a group of adjacent islands.

### Spatial and temporal patterns of threats

Using this database, we firstly described the number of threats impacting threatened species for different spatial scales (globally, on continents, on islands). For our analysis of threat diversity, we used the threats’ first sub-category as the level of threat (e.g., a rail being impacted by sub-categories ‘5.1. Hunting & collecting terrestrial animals’ and ‘5.3. Logging & wood harvesting’ within the threat ‘5. Biological resource use’ was considered as threatened by two threats). Then, we compiled the contemporary threats for all rails, available on the online IUCN database for each species (http://iucnredlist.org). We calculated each threat’s impact score using the *Threat Impact Scoring System* proposed by the IUCN (2019) that defines an overall impact based on scope, severity, and timing and ranges within ‘Negligible’, ‘Low’, ‘Medium’, and ‘High’ impact (https://www.iucnredlist.org/resources/threat-classification-scheme; Fig. S1). We included ‘Past’ impact to illustrate the temporal evolution of threats. The IUCN’s threats classification organises threats in a hierarchical order. At this level, we used the first level of threat (e.g., 1. Residential & commercial development and 2. Agriculture) to be more informative. We split the threat ‘5. Biological resource use’ in ‘Hunting and direct exploitation’ (5.1.) and ‘Logging & indirect effects’ (5.3) to provide relevant conservation policy. If a species had a different impact for threat in 2 sub-categories, we used the higher impact for the classification.

Using this database, we then assessed the spatial pattern of threats for all rails (whether or not they were threatened) by descriptively comparing the type of threats and their impact’s severity i) worldwide, ii) on islands, iii) on continents, and iv) bioregionally (Australasia, Oceania, Nearctic, Neotropics, Palearctic, Afrotropics, Indomalaya; defined following Olson *et al*., 2001). Oceania was grouped with the Australasia bioregion because of its few extant species (five, including three present over the two bioregions). Finally, we compared the contemporary threats with the historical causes of extinctions to assess temporal changes in the threat pattern and distribution of threatened and extinct species.

### Conservation status and gaps

To identify gaps in conservation efforts for rails, we summarised the IUCN’s *Conservation actions classification scheme* and *Research classification scheme* i) globally, ii) per country, and iii) per bioregion, following Olah et al. (2016). Countries and overseas territories were grouped together for country-level analysis. These schemes specify the conservation actions and research needed for each species and were available on the online IUCN database (http://iucnredlist.org). We gathered the possible classifications in five categories: ‘Research, Monitoring and Planning’, ‘Ecosystem Protection and Management’, ‘Species Management’, ‘Education & Awareness’, and ‘Law & Policy’ (i.e., legislative protection). To assess whether conservation status had improved or deteriorated over time, we characterised changes in species’ IUCN conservation status since 1988 (first IUCN Red List assessment) through to 2019 (IUCN, 2019).

### Rails ‘conservation hotspots’ and priority rankings

Rallidae combine species with unique evolutionary traits (e.g., flightlessness) and species with elevated vulnerability (24% of the species are threatened); however to date, there is no framework that accounts for these important aspects in terms of conservation priority. To identify areas with high conservation merit and/or those that deserve improvements in their protection, we conceived a ranking system to classify both world bioregions and countries of high conservation priority for rails (‘conservation hotspots’). Analysis at the country-level grouped countries and their overseas territories together. We created two categories that we deemed useful to reflect important rail conservation: ‘Heritage’ and ‘Threat’. The ‘Heritage’ category aimed to account for species with high conservation value, using endemism and unique evolutionary traits (flightlessness). It was calculated by ranking the number of rail species being either flightless (firstly), island endemic (secondly), or country endemic (thirdly). When relevant, species were classified by one of the above attributes, in this hierarchical order. The ‘Threat’ category was calculated by ranking the number of threatened species (firstly), and the number of species with a worsened IUCN status since their last assessment (secondly).

In cases where ranks were equal, they were split using the country/bioregion’s richness in rail species as the tie breaker (i.e., higher richness would get the higher rank). In this rank system, rank one represents the highest rank. The rank classifications were not made to adequately differentiate between two entities as close scores would not illustrate true difference in conservation priority. We recommend identifying entities with the highest conservations priority as the ones present in both top classifications for the 2 ranks, or as parts of a top 5, top 10, or top 20.

## RESULTS

### Spatial and temporal pattern of threats

All recent rail extinctions (post 1500 A.D.) occurred on islands (n = 26 species: 24 officially extinct and the New Caledonian Rail and Samoan moorhen, considered extinct in this study) and at least 77% were of flightless species (one species was flying and five were not being documented sufficiently to be confident about their flight ability). Of all threatened rail species, the vast majority were island endemics (67%, Table 1, Fig. 2), of which 59% were flightless (39% of all threatened species). The number of threats for threatened island rails (Table 1) was more than twice the known threats that caused extinctions (mean ± SE: 1.6 ± 0.1 threat per extinct species).

**Table 1.** Proportion of threats impacting threatened species at different spatial scale.

| Scale (% threatened sp.) | Median (Q1 – Q3) | Max. | Mean ± SE |
| --- | --- | --- | --- |
| Globally (24%) | 4 (3.0 – 6.0) | 11 | $4.7 \pm 0.4$ |
| Continents (11%) | 4 (4.0 – 8.0) | 11 | $6.2 \pm 0.8$ |
| Islands (52%) | 4 (2.5 – 5.0) | 9 | $4.0 \pm 0.5$ |

**Fig. 2.**
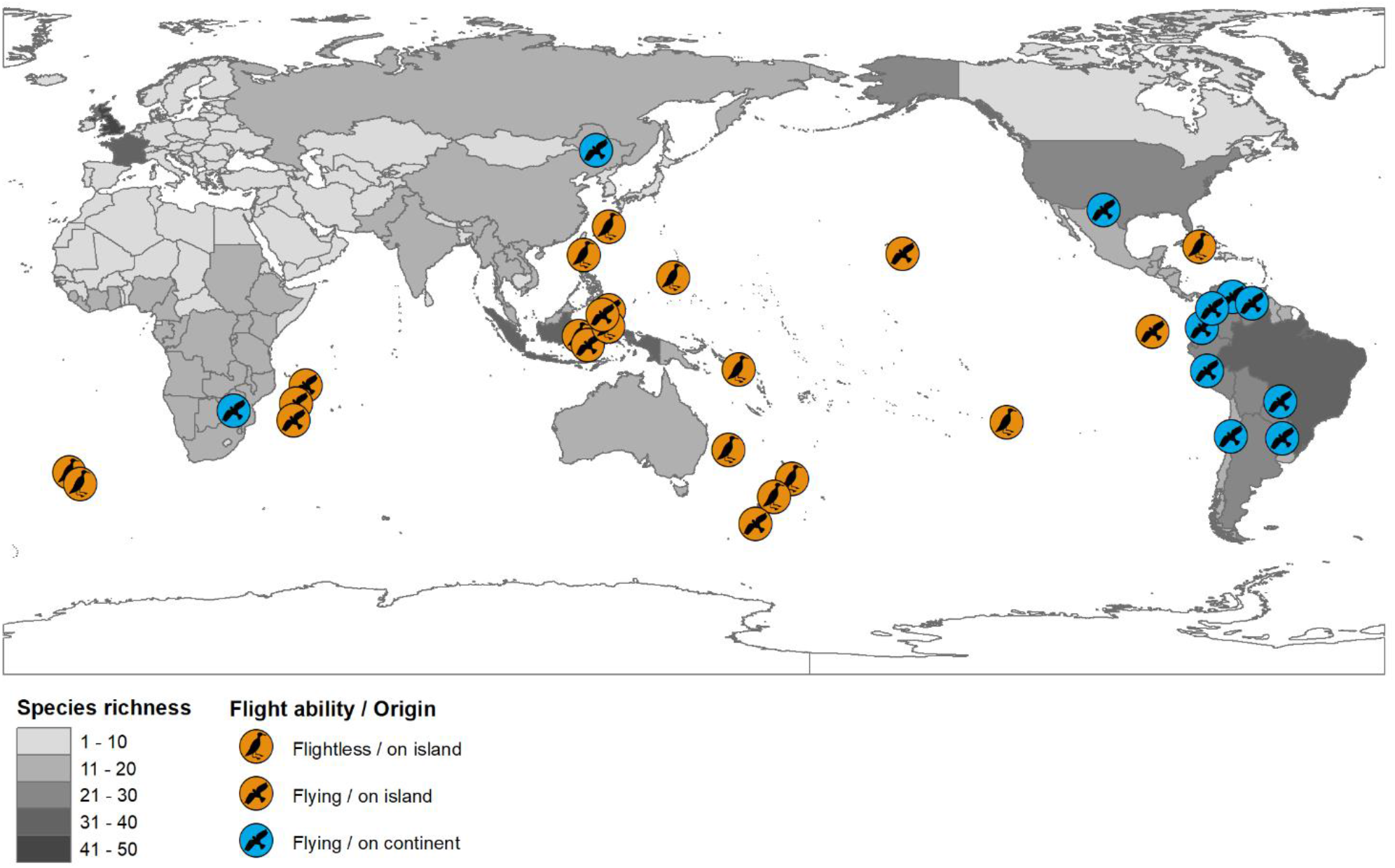
Global distribution of threatened rails, highlighting their origin: continental (blue, n=11), island endemic (orange, n=21) and flying ability (flightless: n=14, 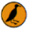). The grey shading represents countries’ total rail richness (including their overseas territories). Projection information: WGS84, centred on 150°E.

Introduced predators and over-hunting were the main drivers to rail extinctions: introduced predators were the only cause in 27% of the cases (partly for 69%) and over-hunting was the only responsible factor in 23% of them (partly for 62%). Habitat loss contributed to 23% of extinctions. Contemporarily, agriculture, invasive species, and logging were the three predominant threats to the extant rails worldwide (impacting 19-26% of species, Fig. 3A). Threats due to agriculture were mostly found in the Afrotropics and the Neotropics, those of invasive species in Australasia/Oceania, and logging in the Afrotropics and Australasia/Oceania (Fig. 3B). For island rails, threats associated with invasive species, hunting, and logging prevailed (38-62% of species), with invasive species (specifically, introduced predators) predominating this group (Fig. 3A). ‘Pollution’ and ‘Natural system modifications’ (e.g., fire, dams, water abstraction) were the only threats that were proportionally more threatening on continents than for island endemics (Fig. 3A). Less than 20% of rails are impacted by ‘Natural System modifications’ globally, however it is an important factor in the Palearctic, where it impacts 60% of that region’s rails (Fig. 3).

**Fig. 3.**
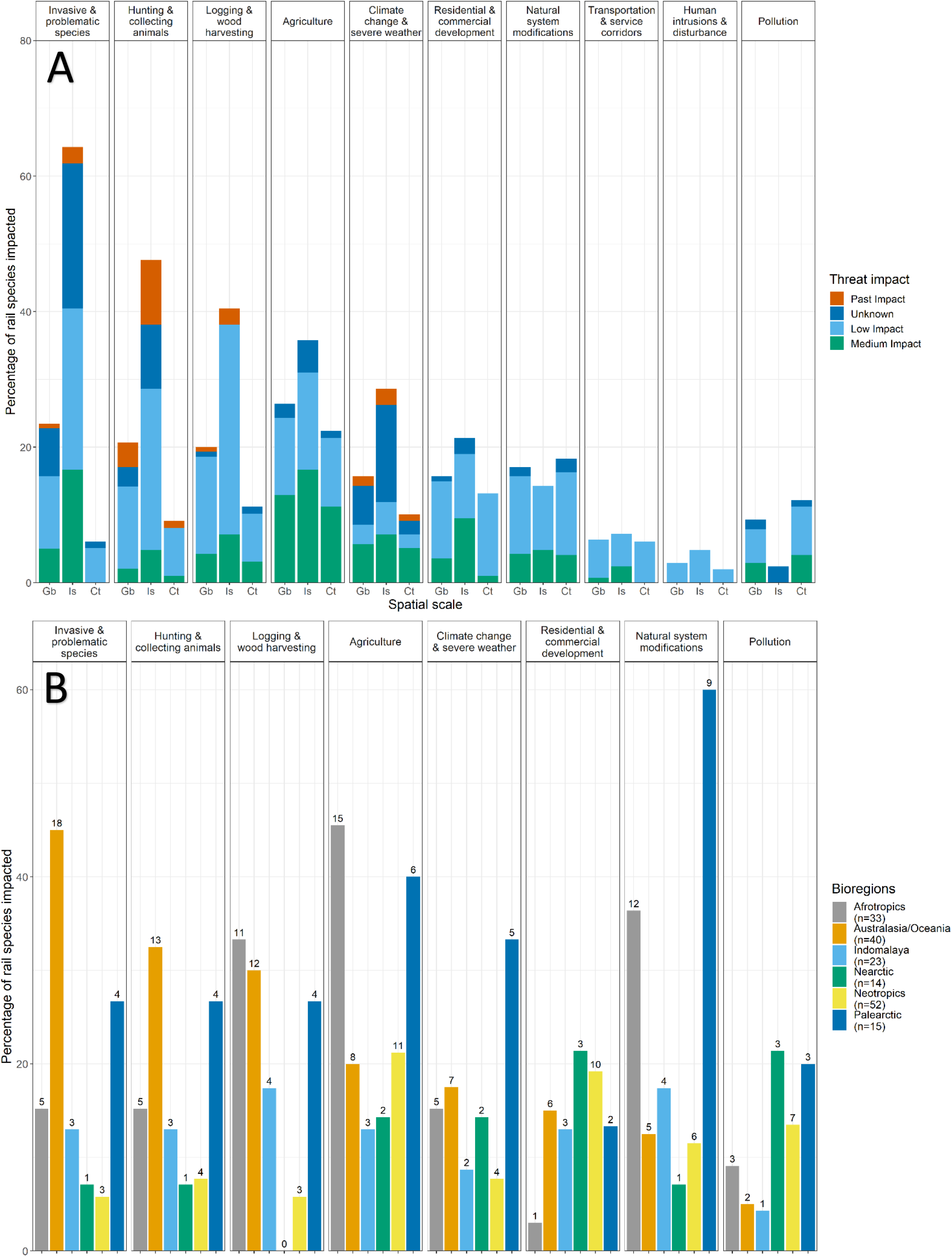
**A** Threat diversity and intensity for rails species globally (Gb; n=140), for island endemics (Is; n=42), and for continental species (Ct; n=98). **B** Threat diversity and percentage of rails impacted for each bioregion. Only threats impacting over 10 species are presented (‘Other’ and ‘Energy production and mining’ are excluded). Bar labels represent the number of species per category.

Most rail extinctions occurred in the Pacific Ocean (65%, Australasia: 8 species, Oceania: 9 species), with 19% in the Indian Ocean (Mascarene islands, 5 species), and 15% on remote Atlantic islands (4 species). Contemporarily, Australasia/Oceania and the Neotropics were the bioregions that host the highest number of threatened rails, representing 35 and 32 species, accounting for 30 and 21% of their total rail species richness, respectively. Indomalaya and Nearctic were the bioregions with the less threatened rails (one species each, respectively *Gallirallus calayanensis* and *Laterallus jamaicensis*).

### Conservation status and gaps

Following the IUCN classification, ‘Research & monitoring’ and ‘Ecosystem protection & management’ were the two most important gaps in conservation efforts directed towards rails, both worldwide (Table 2) and at the country scale (Table S3). They were especially needed in Australasia/Oceania, Palearctic and Nearctic, where up to a quarter of the species required such effort (Table 2). Non-threatened species accounted for 25–79% of the species that required more efforts in ‘Research’ and ‘Ecosystem protection’ categories. Moreover, each threatened species had, on average, 2.5 ± 0.2 conservation actions required, with 45% of them needed more than two types of actions.

**Table 2.** Percentage of species for which conservation gaps were identified by the IUCN Red List (2019), by bioregion. Values in brackets give the proportion of threatened rail species in each conservation gap.

| Bioregion<br>(# species) | Research &<br>Monitoring | Ecosystem<br>protection &<br>management | Species<br>Management | Education &<br>Awareness | Law &<br>Policy |
| --- | --- | --- | --- | --- | --- |
| Australasia/<br>Oceania (40) | 40.0 (22.5) | 27.5 (20.0) | 22.5 (12.5) | 15.0 (5.0) | 7.5 (7.5) |
| Palearctic (15) | 26.7 (13.3) | 20.0 (13.3) | 20.0 (13.3) | 13.3 (6.7) | 13.3 (0.0) |
| Nearctic (14) | 28.6 (7.1) | 28.6 (7.1) | 14.3 (7.1) | 7.1 (0.0) | 0.0 (0.0) |
| Neotropics (52) | 26.9 (21.2) | 23.1 (19.6) | 9.6 (7.7) | 1.9 (1.9) | 0.0 (0.0) |
| Afrotropics (33) | 18.2 (12.1) | 21.2 (12.1) | 6.1 (0.0) | 6.1 (3.0) | 3.0 (0.0) |
| Indomalaya (23) | 13.0 (4.3) | 8.7 (4.3) | 4.3 (4.3) | 4.3 (4.3) | 8.7 (0.0) |
| Globally (140) | 31.7 (20.0) | 27.1 (20.0) | 16.4 (10.7) | 9.3 (5.0) | 4.3 (2.1) |

In the 31-year period between the first Red List assessment (1988) and the most recent (2019), the IUCN conservation status of 11 species worsened, including six species that became newly threatened (Table S4). Eight species were endemic to islands and three were flightless. Conversely, the conservation status improved for seven species, including two downgraded to ‘Least Concern’, one species removed from the threatened category and one species downgraded from ‘Extinct in the Wild’ to ‘Critically Endangered’ thanks to a re-introduction (Table S4).

### Rail ‘conservation hotspots’ and priority rankings

We identified 13 countries present in both top 20 for Heritage and Threat ranks (Table 3). Of them, we could define the five top priority countries (found in both top 10 for Threat and Heritage ranks): Indonesia, New Zealand, the U.S.A., the U.K., and Cuba. The United Kingdom, with its overseas territories included, had the highest rail richness worldwide (33%). All island endemics were found in 14 countries (61% of them were in Australasia/Oceania), and all flightless species in 10 countries (70% of them were in Australasia/Oceania). Of the countries from the top 20 Threat rank, 70% were continental (the U.S.A. and Ecuador were the only continental countries to have island endemics, Table 3). Australasia/Oceania was the most important ‘conservation hotspot’ for rails under our classification system (being top one for ‘Heritage’ and ‘Threat’ ranks, Table S5).

**Table 3.** Classification of countries of highest priority for rail conservation, ordered and ranked by total numbers of: i) threatened, and ii) species with a declined conservation status (Threat rank), iii) flightless, iv) island endemic, and v) country endemic species (Heritage rank). *Ex aequo* countries (equal rank) were split using their total number of species (higher rank for higher richness). Lower values in ranks indicate a higher conservation priority, with rank one representing the highest rank. See Table S6 for details on each category’s rank. Overseas territories are included for each country when relevant. Countries in bold are common between the two ranks.

| Country (#species) | Threat rank | Country (#species) | Heritage rank |
| --- | --- | --- | --- |
| **Indonesia (32)** | 1 | Solomon Islands (9) | 1 |
| **U.S.A. (26)** | 2 | **Indonesia (32)** | 2 |
| Argentina (26) | 3 | Papua New Guinea (16) | 3 |
| **New Zealand (8)** | 4 | **United Kingdom (45)** | 4 |
| **Brazil (31)** | 5 | **New Zealand (8)** | 5 |
| **Madagascar (13)** | 6 | Australia (16) | 6 |
| **United Kingdom (45)** | 7 | **U.S.A. (26)** | 7 |
| Chile (13) | 8 | Philippines (16) | 8 |
| **Cuba (11)** | 9 | **Cuba (11)** | 9 |
| **Peru (27)** | 10 | **Japan (10)** | 10 |
| **Ecuador (23)** | 11 | **Madagascar (13)** | 11 |
| **Colombia (27)** | 12 | **Ecuador (23)** | 12 |
| **Venezuela (21)** | 13 | India (16) | 13 |
| **Japan (10)** | 14 | Seychelles (2) | 14 |
| Ethiopia (15) | 15 | **Venezuela (21)** | 15 |
| Bolivia (26) | 16 | **Brazil (31)** | 16 |
| **Mexico (17)** | 17 | **Colombia (27)** | 17 |
| Zimbabwe (17) | 18 | **Peru (27)** | 18 |
| South Africa (16) | 19 | **Mexico (17)** | 19 |
| Costa Rica (15) | 20 | France (39) | 20 |

## DISCUSSION

### Threat patterns

While most threatened bird species are found on continents (Bennett *et al*., 2001), the rails show a different pattern. All extinct and most threatened rail species are island endemics, including most flightless species. Most of the past rail extinctions occurred on the Pacific islands, directly linked to the fact that they were the support for the largest radiation of rails, reaching high levels of endemism. Flightlessness is an evolutionary trait that has been found to make bird species more extinction-prone during different waves of extinction, in general (Duncan *et al*., 2002; Boyer, 2008) and specifically in rails (Steadman, 1995; Curnutt & Pimm, 2001), and also appears as an important contributor to the vulnerability of rails to contemporary threatening processes.

From the three main threats recognised from past extinctions (over-hunting, introduced predators, and habitat modification), anthropogenic activities have diversified to impact rails with up to eleven different threats, as recognised by the IUCN, with some of these threats affecting up to 60% of all island rails. The three key threats that we identified for extant rails – impacting about a fifth of all rails globally – are agriculture, invasive species, and logging, a pattern that is consistent with other bird species globally (BirdLife International, 2017). If we also include hunting, these four are also the most impactful threats for island endemic rails (with a different order of impact, and invasive predators having the most threatening consequences). Nevertheless, invasive species are not the main concern for other threatened-island-bird species globally, where over-exploitation and agriculture predominate (Leclerc *et al*., 2018). The overwhelming threat posed by invasive predators to island rails could be linked to vulnerable adaptations like flightlessness, predator naivete (Duncan *et al*., 2002; Steadman, 2006; Boyer, 2008) and their territorial lifestyle (e.g., ground-nesting), but more research is needed to disentangle rails’ vulnerability in depth.

Here, we have demonstrated that one threatening pathway has not changed between extinct and threatened island rails: invasive predators are consistently the key problem (extinct sp.: 96%, threatened sp.: 62%), followed by over-hunting (extinct sp.: 54%, threatened sp.: 38%). Australasia/Oceania, where the problem of invasive predators is most prevalent, also has the most extinct and threatened rail species, and hosts 61% of the diversity of island endemic rails and 70% of all surviving flightless rails, making it the top one bioregion for conservation priority. However, Oceania, which in ancient times supported hundreds of flightless rails, now stands bleakly as a rail-species graveyard after the Holocene mass extinction (only five rail species survive in the Pacific basin, including just two endemics to the region).

On the other hand, ‘Agriculture’, ‘Natural system modifications’ (e.g., fire, dams, water abstraction), and ‘Residential and commercial development’ are the prevailing threats to continental rails, making the Neotropics the top 2 conservation priority bioregion. Compared to island rails, we found that continental rails were impacted by a wider diversity of threats. This difference could be due to continental rails being intrinsically more resistant to threats, even in synergy, or because fewer threats might naturally occur on islands. To date, no rail species has gone extinct on continents at human contact, but escalating contemporary threats could be creating another pathway in the threatening process to rails that could act to undermine their populations’ resilience and so jeopardise the persistence of continental species in the long-term, especially if the threats continue to intensify or act in synergy (Côté *et al*., 2016; Leclerc *et al*., 2018). Only 11% of the continental rails are threatened, nevertheless, the IUCN Red List focussing on particular regions (such as the European Red List; https://www.iucnredlist.org/regions/europe) illustrates that some of the ‘Least Concern’ continental rails can appear as threatened at a smaller scale and therefore, could be overlooked for conservation.

### Conservation efforts, gaps, and priorities

We found ‘Research and monitoring’ effort to be the most important gap in rail conservation globally (25–42% of species in half the bioregions), with ‘Ecosystem protection and management’ almost as equally lacking (25–29% in half the bioregions). Likewise, de Lima *et al*. (2011) found that the degree of research effort was biased across all bird species and areas. Moreover, threatened bird species tend to be less well-researched than non-threatened ones, probably due to their smaller geographical range (Brooks *et al*., 2008). Indeed, the Neotropics and Australasia/Oceania host the greatest number of threatened rail species and of threatened bird species in general, but both are understudied (0.03 and 0.08 published papers per threatened bird species, respectively; Brooks *et al*., 2008).

By analysing the threat patterns across spatial scales, we uncovered important regional variation in the ranking of threats to rails (e.g., ‘Natural System modifications’ impacts 60% of the Palearctic species, compared to only 17% globally), allowing us to address conservation more specifically. Our analysis also revealed a positive outcome, as seven species improved their conservation status recently. However, the number of species whose IUCN status is deteriorating is increasing faster than the ones improving. Hence, it appears as equally important to work on improving knowledge and protection for overlooked species, and especially for the two ‘Data Deficient’ rails, the Brown-banded Rail (*Lewinia mirifica*) and the Colombian Crake (*Neocrex colombiana*).

While Brooks *et al*. (2008) found that most threatened bird species inhabit low-income countries, the top countries in our threat rank for rails (the most threatened species and species with a declining IUCN status) include a mix of low- and high-income countries, with New Zealand, and U.S.A. within in the top four. Low-income countries can be expected to carry threatened species as they typically have less enforcement and higher rates of wildlife poaching and habitat loss (Kerr & Currie, 1995; Green, 1996; Blaikie & Jeanrenaud, 1997; Olah *et al*., 2016). However, in the case of rails, high-income countries have more threatened species because they possess, by chance, many islands supporting island endemic rails that are endangered due to different threatening mechanisms, such as invasive predators, that are not directly linked to a countries’ income.

A few countries identified as rail ‘conservation hotspots’ (i.e., with high number of threatened rails and with high-conservation-value species) were also the ones most lacking in conservation efforts. In that regard, we suggest that Indonesia, the U.S.A., the United Kingdom, New Zealand, and Cuba should be focusing more on measures to protect and recover their rail species, equally on their mainland and overseas territories. The Solomon Islands is the country hosting the most flightless rails (top 2 ‘conservation hotspot’, Table S6), and as only 19 flightless rails remain globally compared to the many hundreds existing in the Holocene (Steadman, 1995; Curnutt & Pimm, 2001), they should also be given particularly directed conservation attention.

Climate change also looms as a future threat for all island birds (Bellard et al., 2014), especially via its impact on sea-level rise (Curnutt & Pimm, 2001; Garnett & Reside, 2018) and habitat loss through ecosystem degradation (IUCN, 2019). This would be particularly relevant in the Pacific region, that risks seeing the remaining island endemics wiped out under the impacts of climate change. As already found for the Hawaiian common gallinule (*Gallinula galeata sandvicensis*; van Rees & Reed, 2018), climate change’s impacts can reduce habitat quality (and therefore an island’s already limited carrying capacity), which can be especially threatening for rails endemic to small oceanic isolates — e.g. the Guam Rail (*Hypotaenidia owstoni*), the Inaccessible Island rail (*Atlantisia rogersi*) and the Lord Howe woodhen (*Gallirallus sylvestris*). Further research is needed to forecast the impacts of climate change on such restricted species in order to accurately determine the most adequate conservation measures to implement, such as defining climatic refugia, translocations or captivity plans (Seddon *et al*., 2014; Braidwood *et al*., 2018; Garnett & Reside, 2018; Beaumont *et al*., 2019).

### Conclusion

The threat pattern in rails follows two different pathways. The world’s extant island rails are continuing on the same trajectory as extinct species, with the majority of island endemics still suffering from the same key threats that drove others extinct (invasive predators and over-hunting). However, the synergy of modern threats has also created another trajectory, whereby continental rails are impacted by a diversity of contemporary anthropogenic activities that could jeopardise the long-term survival of previously resilient populations. This dichotomy leads to a complex pattern under which conservation options are branching into various profiles, whose priority will depend on the type of rail species (continental or island endemic) and the relevant scale (bioregions, countries, islands). This synthesis provides the first analysis of the spatio-temporal threat pattern in rails, and further research is needed to disentangle the role of extrinsic and intrinsic traits in threatening mechanisms, and thus better anticipate future rail extinctions.

## Acknowledgements

We thank Stefania Ondei for her help with ArcGIS and all individuals and organisations that participated to bird assessments for the IUCN RedList.

## REFERENCES

Beaumont, L.J., Esperón-Rodríguez, M., Nipperess, D.A., Wauchope-Drumm, M. & Baumgartner, J.B. (2019) Incorporating future climate uncertainty into the identification of climate change refugia for threatened species. Biological Conservation, 237, 230–237.

Bellard, C., Leclerc, C. & Courchamp, F. (2014) Impact of sea level rise on the 10 insular biodiversity hotspots. Global Ecology and Biogeography, 23, 203–212.

Bennett, P.M., Owens, I.P.F. & Baillie, J.E.M. (2001) The History and Ecological Basis of Extinction and Speciation in Birds. Biotic Homogenization (ed. by J.L. Lockwood and M.L. Mckinney), pp. 201–222. Springer US, Boston, MA.

BirdLife International (2017) A range of threats drives declines in bird populations. Downloaded from http://www.birdlife.org_on_15/07/2019.

BirdLife International (2018) <http://datazone.birdlife.org/home>, accessed on 21/02/2018.

Blaikie, P. & Jeanrenaud, S. (1997) Biodiversity and human welfare. Social change and conservation. Environmental politics and impacts of national parks and protected areas, 46–70.

Boyer, A.G. (2008) Extinction patterns in the avifauna of the Hawaiian islands. Diversity and Distributions, 14, 509–517.

Braidwood, D.W., Taggart, M.A., Smith, M. & Andersen, R. (2018) Translocations, conservation, and climate change: use of restoration sites as protorefuges and protorefugia. Restoration Ecology, 26, 20–28.

Brooks, T.M., Collar, N.J., Green, R.E., Marsden, S.J. & Pain, D.J. (2008) The science of bird conservation. Bird Conservation International, 18, S2–S12.

Côté, I.M., Darling, E.S. & Brown, C.J. (2016) Interactions among ecosystem stressors and their importance in conservation. Proceedings of the Royal Society B: Biological Sciences, 283, 20152592.

Croxall, J.P., Butchart, S.H., Lascelles, B., Stattersfield, A.J., Sullivan, B., Symes, A. & Taylor, P. (2012) Seabird conservation status, threats and priority actions: a global assessment. Bird Conservation International, 22, 1–34.

Curnutt, J. & Pimm, S. (2001) How many bird species in Hawaii and the Central Pacific before first contact? Studies in Avian Biology, 22, 15–30.

de Lima, R.F., Bird, J.P. & Barlow, J. (2011) Research effort allocation and the conservation of restricted-range island bird species. Biological Conservation, 144, 627–632.

Dias, M.P., Martin, R., Pearmain, E.J., Burfield, I.J., Small, C., Phillips, R.A., Yates, O., Lascelles, B., Borboroglu, P.G. & Croxall, J.P. (2019) Threats to seabirds: A global assessment. Biological Conservation, 237, 525–537.

Duncan, R.P., Blackburn, T.M. & Worthy, T.H. (2002) Prehistoric bird extinctions and human hunting. Proceedings of the Royal Society of London. Series B: Biological Sciences, 269, 517–521.

Garcia-R, J.C., Lemmon, E.M., Lemmon, A.R. & French, N. (2020) Phylogenomic Reconstruction Sheds Light on New Relationships and Timescale of Rails (Aves: Rallidae) Evolution. Diversity, 12, 70.

Garcia-R., J.C., Gibb, G.C. & Trewick, S.A. (2014) Deep global evolutionary radiation in birds: diversification and trait evolution in the cosmopolitan bird family Rallidae. Molecular Phylogenetics and Evolution, 81, 96–108.

Garnett, S.T. & Reside, A.E. (2018) Managing island in the context of climate change. Australian Island Arks: Conservation, Management and Opportunities. Ed. by Dorian Moro, Derek Ball, Sally Bryant.

Green, A.J. (1996) Analyses of Globally Threatened Anatidae in Relation to Threats, Distribution, Migration Patterns, and Habitat Use. Conservation Biology, 10, 1435–1445.

IPBES (2019) (Intergovernmental Science-Policy Platform on Biodiversity and Ecosystem Services). “Nature’s dangerous decline ‘unprecedented,’ species extinction rates ‘accelerating’.” ScienceDaily, 6 May 2019. <https://www.sciencedaily.com/releases/2019/05/190506093610.htm>

IUCN (2019) The IUCN Red List of Threatened Species. Version 2019-3. Available at: http://www.iucnredlist.org (accessed 12 December 2019)

Kerr, J.T. & Currie, D.J. (1995) Effects of Human Activity on Global Extinction Risk. Conservation Biology, 9, 1528–1538.

Leclerc, C., Courchamp, F. & Bellard, C. (2018) Insular threat associations within taxa worldwide. Scientific reports, 8, 6393.

Olah, G., Butchart, S.H.M., Symes, A., Guzmán, I.M., Cunningham, R., Brightsmith, D.J. & Heinsohn, R. (2016) Ecological and socio-economic factors affecting extinction risk in parrots. Biodiversity and Conservation, 25, 205–223.

Olson, D.M., Dinerstein, E., Wikramanayake, E.D., Burgess, N.D., Powell, G.V., Underwood, E.C., D’amico, J.A., Itoua, I., Strand, H.E. & Morrison, J.C. (2001) Terrestrial Ecoregions of the World: A New Map of Life on Earth. A new global map of terrestrial ecoregions provides an innovative tool for conserving biodiversity. BioScience, 51, 933–938.

Owens, I.P.F. & Bennett, P.M. (2000) Ecological basis of extinction risk in birds: Habitat loss versus human persecution and introduced predators. Proceedings of the National Academy of Sciences, 97, 12144–12148.

Rosenberg, K.V., Dokter, A.M., Blancher, P.J., Sauer, J.R., Smith, A.C., Smith, P.A., Stanton, J.C., Panjabi, A., Helft, L., Parr, M. & Marra, P.P. (2019) Decline of the North American avifauna. Science, 366, 120–124.

Seddon, P.J., Griffiths, C.J., Soorae, P.S. & Armstrong, D.P. (2014) Reversing defaunation: restoring species in a changing world. Science, 345, 406–412.

Spatz, D.R., Newton, K.M., Heinz, R., Tershy, B., Holmes, N.D., Butchart, S.H.M. & Croll, D.A. (2014) The Biogeography of Globally Threatened Seabirds and Island Conservation Opportunities. Conservation Biology, 28, 1282–1290.

Steadman, D.W. (1995) Prehistoric extinctions of Pacific island birds: Biodiversity meets zooarchaeology. Science (New York, N.Y.), 267, 1123.

Steadman, D.W. (2006) Extinction and biogeography of tropical Pacific birds. University of Chicago Press.

Taylor, B. & van Perlo, B. (1998) Rails: a guide to rails, crakes, gallinules and coots of the world. Pica Press, Mountfield, U.K.

van Rees, C.B. & Reed, J.M. (2018) Predicted effects of landscape change, sea level rise, and habitat management on the extirpation risk of the Hawaiian common gallinule (Gallinula galeata sandvicensis) on the island of O’ahu. PeerJ, 6, 4990.

