## Supplementary material for "Characterising the spatio-temporal threats, conservation hotspots, and conservation gaps for the most extinction-prone bird family (Aves: Rallidae)"

**Table S1** Classification of countries of highest priority for rail conservation, excluding the rallid sub-family of Sarothruridae, ordered and ranked by total numbers of: i) threatened, and ii) species with a declined conservation status (Threat rank), iii) flightless, iv) island endemic, and v) country endemic species (Heritage rank). *Ex aequo* countries (equal rank) were split using their total number of species (higher rank for higher richness). Lower values in ranks indicate a higher conservation priority, with rank one representing the highest rank. Overseas territories are included for each country when relevant. Countries in bold are common between the two ranks.

| Country (#species) | Threat rank | Country (#species) | Heritage rank |
| --- | --- | --- | --- |
| **Indonesia (28)** | 1 | Solomon Islands (9) | 1 |
| **U.S.A. (26)** | 2 | **Indonesia (28)** | 2 |
| Argentina (26) | 3 | Papua New Guinea (13) | 3 |
| **New Zealand (8)** | 4 | **United Kingdom (45)** | 4 |
| **Brazil (31)** | 5 | **New Zealand (8)** | 5 |
| **United Kingdom (45)** | 6 | Australia (16) | 6 |
| Chile (13) | 7 | **U.S.A. (26)** | 7 |
| **Cuba (11)** | 8 | Philippines (16) | 8 |
| **Peru (27)** | 9 | **Cuba (11)** | 9 |
| **Ecuador (23)** | 10 | **Japan (10)** | 10 |
| **Madagascar (9)** | 11 | **Madagascar (9)** | 11 |
| **Colombia (27)** | 12 | **Ecuador (23)** | 12 |
| **Venezuela (21)** | 13 | India (16) | 13 |
| **Japan (10)** | 14 | Seychelles (2) | 14 |
| Bolivia (26) | 15 | **Venezuela (21)** | 15 |
| **Mexico (17)** | 16 | **Brazil (31)** | 16 |
| Costa Rica (15) | 17 | **Colombia (27)** | 17 |
| Panama (14) | 18 | **Peru (27)** | 18 |
| Belize (13) | 19 | **Mexico (17)** | 19 |
| Honduras (11) | 20 | France (39) | 20 |

**Table S2.** List of rail species excluded from the analyses on extant species.

| Common name | Latin name | IUCN |
| --- | --- | --- |
| Brown-banded Rail | *Lewinia mirifica* | Data Deficient |
| Colombian Crake | *Neocrex colombiana* | Data Deficient |
| New Caledonian rail | *Gallirallus lafresnayanus* | Critically endangered* |
| Samoan moorhen | *Pareudiastes pacificus* | Critically endangered* |

*****have not been seen with certainty since the 19^th^ century and are suspected to be extinct.

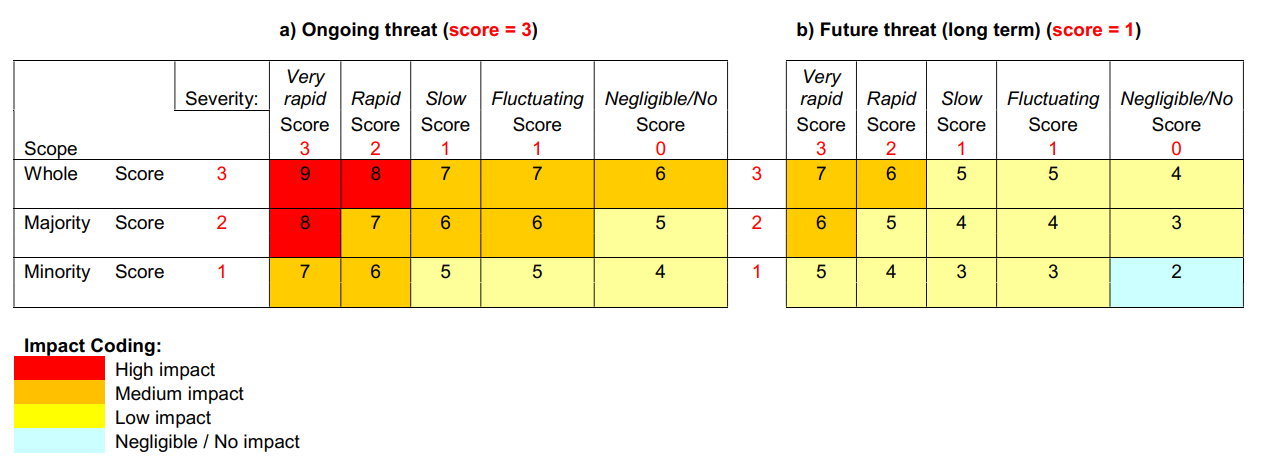
**Fig. S1.** IUCN’s *Threat impact scoring system* (based on additive scores and defined thresholds). Version 1.0. Accessible at <https://nc.iucnredlist.org/redlist/content/attachment_files/Dec_2012_Guidance_on_Threat_Impact_Scoring_Revised.pdf>

**Table S3** Number of conservation gaps identified by the IUCN Red List (2019) per country (including their overseas territories). Only countries with over five conservation gaps are presented (100 countries have five or less gaps, and 84 countries where rails are present have no gap identified). Values between brackets are the subset of rail species with conservation gaps that are threatened species. Countries include their overseas territories.

| Country (# species) | Research & Monitoring | Ecosystem protection & management | Species Management | Education & Awareness | Law & Policy | Total gaps |
| --- | --- | --- | --- | --- | --- | --- |
| Indonesia (32) | 9 (4) | 5 (3) | 2 (0) | 3 (1) | 1 (0) | 20 |
| United Kingdom (45) | 2 (1) | 4 (3) | 4 (3) | 2 (1) | 1 (0) | 13 |
| U.S.A. (26) | 4 (2) | 4 (2) | 3 (2) | 1 (0) | 1 (1) | 13 |
| Mexico (17) | 4 (1) | 4 (1) | 2 (1) | 1 (0) | 0 (0) | 11 |
| Argentina (26) | 4 (3) | 3 (3) | 2 (1) | 1 (1) | 0 (0) | 10 |
| China (18) | 3 (1) | 2 (1) | 2 (1) | 1 (0) | 2 (0) | 10 |
| PNG (16) | 4 (0) | 2 (0) | 2 (0) | 2 (0) | 0 (0) | 10 |
| Russia (13) | 3 (1) | 2 (1) | 2 (1) | 1 (0) | 2 (0) | 10 |
| Ethiopia (15) | 2 (1) | 3 (1) | 2 (0) | 1 (0) | 1 (0) | 9 |
| Madagascar (13) | 4 (3) | 4 (3) | 0 (0) | 1 (1) | 0 (0) | 9 |
| New Zealand (8) | 3 (3) | 2 (2) | 3 (3) | 1 (1) | 0 (0) | 9 |
| Chile (13) | 3 (2) | 2 (2) | 2 (1) | 1 (1) | 0 (0) | 8 |
| Solomon Islands (9) | 3 (1) | 1 (1) | 1 (0) | 2 (0) | 1 (1) | 8 |
| Mongolia (6) | 2 (1) | 2 (1) | 2 (1) | 1 (0) | 1 (0) | 8 |
| Brazil (31) | 3 (3) | 3 (3) | 1 (1) | 0 (0) | 0 (0) | 7 |
| Zimbabwe (17) | 2 (1) | 2 (1) | 1 (0) | 1 (0) | 1 (0) | 7 |
| Philippines (16) | 2 (1) | 1 (1) | 2 (1) | 2 (1) | 0 (0) | 7 |
| South Africa (16) | 2 (1) | 2 (1) | 1 (0) | 1 (0) | 1 (0) | 7 |
| Cuba (11) | 3 (2) | 3 (2) | 1 (1) | 0 (0) | 0 (0) | 7 |
| Japan (10) | 2 (2) | 2 (2) | 2 (2) | 1 (1) | 0 (0) | 7 |

Note: When excluding the family of Sarothruridae, Madagascar, Ethiopia, and Zimbabwe were excluded from the rank.

**Table S4.** Rails species that have worsened or improved their latest IUCN conservation status.

| Latin name | Current status | Previous status | Direction of change | Date of change | Flightless -ness |
| --- | --- | --- | --- | --- | --- |
| *Cyanolimnas cerverai* | CR | EN | Worsened | 2010 | Flightless |
| *Fulica alai* | VU | Unknown (LR/LC) | Worsened | 1994 | Flying |
| *Gallirallus australis* | VU | Unknown (LR/LC) | Worsened | 2000 | Flightless |
| *Hypotaenidia rovianae* | NT | Unknown (LR/LC) | Worsened | 2004 | Flightless |
| *Laterallus jamaicensis* | EN | NT | Worsened | 2019 | Flying |
| *Laterallus spilonota* | VU | NT | Worsened | 2000 | Flying |
| *Lewinia muelleri* | VU | Unknown (LR/LC) | Worsened | 1994 | Flying |
| *Rallicula leucospila* | NT | LC | Worsened | 2004 | Flying |
| *Rallus madagascariensis* | VU | Unknown (LR/LC) | Worsened | 2004 | Flying |
| *Rougetius rougetii* | NT | Unknown (LR/LC) | Worsened | 2004 | Flying |
| *Sarothrura ayresi* | CR | EN | Worsened | 2013 | Flying |
| *Fulica cornuta* | NT | Unknown (LR/NT) | Improved | 2004 | Flying |
| *Hypotaenidia owstoni* | CR | EW | Improved | 2019 | Flightless |
| *Laterallus levraudi* | VU | EN | Improved | 2018 | Flying |
| *Megacrex inepta* | LC | NT | Improved | 2017 | Flightless |
| *Rallina canningi* | LC | NT | Improved | 2017 | Flying |
| *Rallus antarcticus* | VU | CR | Improved | 2000 | Flying |
| *Zapornia olivieri* | EN | CR | Improved | 2004 | Flying |

**Table S5**. Classification of bioregions of highest priority for rail conservation, ordered and ranked by total numbers of rail: i) threatened, and ii) species with a declined conservation status (Threat rank), iii) flightless, iv) island endemic, and v) country endemic species (Heritage rank). *Ex aequo* bioregions (equal rank) were split using their total number of species (higher rank for higher richness). Lower values in ranks indicate a higher priority, with rank one representing the highest rank. Bioregion ‘Other’ indicates Inaccessible, Gough, and Henderson islands and was not included in the classification.

| Bioregion (# species) | Threat-ened | Degraded status | Threat rank | Flight-less | Island endemic | Country endemic | Heritage rank |
| --- | --- | --- | --- | --- | --- | --- | --- |
| Australasia/Oceania (40) | 12 | 5 | 1 | 14 | 13 | 3 | 1 |
| Neotropics (52) | 11 | 4 | 2 | 1 | 1 | 6 | 3 |
| Afrotropics (33) | 4 | 3 | 3 | 0 | 7 | 0 | 5 |
| Palearctic (15) | 2 | 0 | 4 | 1 | 0 | 0 | 4 |
| Nearctic (14) | 1 | 1 | 5 | 0 | 0 | 1 | 6 |
| Indomalaya (23) | 1 | 0 | 6 | 1 | 3 | 0 | 2 |
| Other (3) | 3 | 0 | / | 3 | 0 | 0 | / |

**Table S6**. Classification of countries of highest priority for rail conservation, ordered and ranked by total numbers of rail: i) threatened, and ii) with a declined conservation status (Threat rank), iii) flightless, iv) island endemic, and v) country endemic species (Heritage rank). *Ex aequo* countries (equal rank) were split using their total number of species (higher rank for higher richness). Lower values in ranks indicate a higher priority, with rank one representing the highest rank. Countries include their overseas territories.

| Country (#species) | Threat  -ened | Degraded status | Threat rank | Flight-less | Island endemic | Country endemic | Heritage rank |
| --- | --- | --- | --- | --- | --- | --- | --- |
| Indonesia (32) | 5 | 1 | 1 | 3 | 10 | 0 | 2 |
| U.S.A. (26) | 3 | 2 | 2 | 1 | 2 | 1 | 7 |
| Argentina (26) | 3 | 2 | 3 | 0 | 0 | 0 | 21 |
| New Zealand (8) | 3 | 2 | 4 | 2 | 1 | 0 | 5 |
| Brazil (31) | 3 | 1 | 5 | 0 | 0 | 1 | 16 |
| Madagascar (13) | 3 | 1 | 6 | 0 | 7 | 0 | 11 |
| U.K. (45) | 3 | 0 | 7 | 3 | 0 | 0 | 4 |
| Chile (13) | 2 | 2 | 8 | 0 | 0 | 0 | 49 |
| Cuba (11) | 2 | 2 | 9 | 1 | 0 | 0 | 9 |
| Peru (27) | 2 | 1 | 10 | 0 | 0 | 1 | 18 |
| Ecuador (23) | 2 | 1 | 11 | 0 | 1 | 0 | 12 |
| Colombia (27) | 2 | 0 | 12 | 0 | 0 | 1 | 17 |
| Venezuela (21) | 2 | 0 | 13 | 0 | 0 | 2 | 15 |
| Japan (10) | 2 | 0 | 14 | 1 | 0 | 0 | 10 |
| Ethiopia (15) | 1 | 2 | 15 | 0 | 0 | 0 | 36 |
| Bolivia (26) | 1 | 1 | 16 | 0 | 0 | 0 | 22 |
| Mexico (17) | 1 | 1 | 17 | 0 | 0 | 1 | 19 |
| Zimbabwe (17) | 1 | 1 | 18 | 0 | 0 | 0 | 30 |
| South Africa (16) | 1 | 1 | 19 | 0 | 0 | 0 | 33 |
| Costa Rica (15) | 1 | 1 | 20 | 0 | 0 | 0 | 35 |
| Solomon Is. (9) | 1 | 1 | 24 | 4 | 0 | 0 | 1 |
| Australia (16) | 1 | 0 | 31 | 2 | 0 | 2 | 6 |
| Philippines (16) | 1 | 0 | 32 | 1 | 2 | 0 | 8 |
| France (39) | 0 | 0 | 39 | 0 | 0 | 0 | 20 |
| Papua New Guinea (16) | 0 | 0 | 45 | 3 | 4 | 0 | 3 |
| India (16) | 0 | 0 | 46 | 0 | 1 | 0 | 13 |
| Seychelles (2) | 0 | 0 | 188 | 0 | 1 | 0 | 14 |
